# The actin binding protein profilin 1 is critical for mitochondria function

**DOI:** 10.1101/2023.08.07.552354

**Authors:** Tracy-Ann Read, Bruno A. Cisterna, Kristen Skruber, Samah Ahmadieh, Halli L. Lindamood, Josefine A. Vitriol, Yang Shi, Austin E.Y.T. Lefebvre, Joseph B. Black, Mitchell T. Butler, James E. Bear, Alena Cherezova, Daria V. Ilatovskaya, Neil L. Weintraub, Eric A. Vitriol

## Abstract

Profilin 1 (PFN1) is an actin binding protein that is vital for the polymerization of monomeric actin into filaments. Here we screened knockout cells for novel functions of PFN1 and discovered that mitophagy, a type of selective autophagy that removes defective or damaged mitochondria from the cell, was significantly upregulated in the absence of PFN1. Despite successful autophagosome formation and fusion with the lysosome, and activation of additional mitochondrial quality control pathways, PFN1 knockout cells still accumulate damaged, dysfunctional mitochondria. Subsequent imaging and functional assays showed that loss of PFN1 significantly affects mitochondria morphology, dynamics, and respiration. Further experiments revealed that PFN1 is located to the mitochondria matrix and is likely regulating mitochondria function from within rather than through polymerizing actin at the mitochondria surface. Finally, PFN1 mutants associated with amyotrophic lateral sclerosis (ALS) fail to rescue PFN1 knockout mitochondrial phenotypes and form aggregates within mitochondria, further perturbing them. Together, these results suggest a novel function for PFN1 in regulating mitochondria and identify a potential pathogenic mechanism of ALS-linked PFN1 variants.

## Introduction

The actin cytoskeleton is a dynamic structural network within eukaryotic cells which regulates their shape and behavior. It is indispensable for driving cellular processes including motility, cytokinesis, and intracellular communication^1-3^. Inside the cell, forces generated by the assembly and organization of actin filaments are used to control the dynamics of organelles like mitochondria^4,5^. Proper mitochondrial function requires precise control over the shape and distribution of mitochondria networks, which is achieved through mitochondria fission, fusion, transport, and clearance of damaged organelles^6-8^. Actin has been implicated in all these regulatory steps, largely through the polymerization of filaments on the mitochondria surface or contact sites on the endoplasmic mitochondria transport and physiology through reticulum^9,10^. For example, INF2/Spire1C and Arp2/3-mediated filament assembly are essential components of the fission process^4,5^. Moreover, tethering of mitochondria to actin filaments via Miro or Myosin 19 can alter morphological changes^8,11^.

Actin filaments are also crucial for the activation of metabolic pathways that in turn regulate mitochondrial function. For instance, activation of the glycolytic enzymes’ aldolase and glyceraldehyde phosphate dehydrogenase, can occur by direct binding to F-actin^12,13^. Furthermore, TRIM21 (Tripartite Motif-containing Protein 21) is sequestered by F-actin bundles, thereby reducing its access to substrates such as the rate-limiting metabolic enzyme phosphofructokinase (PFK), thus ensuring high glycolytic rates^14^. In neurons, actin in mitochondria is needed for the retention of cytochrome c between respiratory chain complexes III and IV through direct association with both complexes^15^. Additionally, actin depolymerization with cytochalasin b can enhance mitochondrial respiration through increased complex IV activity^15^.

Here, we investigate the role of profilin 1 (PFN1) in controlling mitochondria function. PFN1 is an actin monomer-binding protein that has profound influence over actin filament assembly. Profilin prevents spontaneous filament assembly, promotes the exchange of ADP-actin to the polymerization competent ATP-bound form, and even controls which filaments assemble from the monomer pool through its interactions with different actin polymerases^3,16^. However, apart from mitochondria dysfunction associated with amyotrophic lateral sclerosis (ALS) associated mutants of PFN1^17-19^, there is currently no evidence to support a direct link between PFN1 and mitochondria. Here, we deleted the *PFN1* gene in two different cell lines and show that PFN1 is critical for mitochondria function. Loss of PFN1 causes a massive upregulation of quality control pathways due to the accumulation of damaged mitochondria. We show that PFN1 is important for mitochondria respiration, morphology, and dynamics, and that ALS-associated mutants fail to rescue mitochondria dysfunction caused by PFN1 loss-of-function. Most surprisingly, we provide compelling evidence that PFN1 localizes not to the outside of mitochondria, but to the inside of the mitochondrial matrix. ALS-associated PFN1 mutants can also aggregate inside of mitochondria, causing even more damage. Thus, not only is PFN1 a critical regulator of mitochondria function, but we have identified a causative role for mutant PFN1 in cellular toxicity through combined loss- and gain-of-toxic function effects.

## Results

### Loss of PFN1 causes an upregulation of autophagy through mTOR deactivation

In previous work we used CRISPR/Cas9 to knock out (KO) PFN1 in the central nervous system-derived cath.A differentiated (CAD) cell line^20^. Interestingly, RNA-seq analysis identified significant changes in expression of genes associated with lysosome/endosome systems and autophagy^21^ upon the loss of PFN1 expression (Fig. 1A). This included upregulation of genes in the lysosomal proteolytic system (*CTSZ, CTSA)*, genes in the endosome/lysosome pathway (*TPCN2, UNC93B1, VPS16, VPS9*), and autophagy activating genes such as *Atg13, USP11, HDAC10 and PRKDC*. Furthermore, PFN1 KO cells had differential expression of mammalian target of rapamycin (mTOR) pathway deactivating genes ^21^(*GRB10, DEPTOR, TSC1, RRAGD*) (Fig. 1A), which is the predominant signaling event that induces autophagy. We confirmed the upregulation of *DEPTOR* (Fig. 1B) and a decrease in phosphorylated mTOR (Fig. 1C) in PFN1 KO cells via quantitative western blotting. We also expressed mRuby2-LC3, which forms puncta when autophagosomes are formed,^22,23^ and found a three-fold increase in LC3 puncta in PFN1 KO cells. Importantly, this phenotype could be rescued by expressing a GFP-PFN1 construct (Fig. 1D). Thus, autophagy is significantly upregulated in the absence of PFN1.

**Figure 1.**
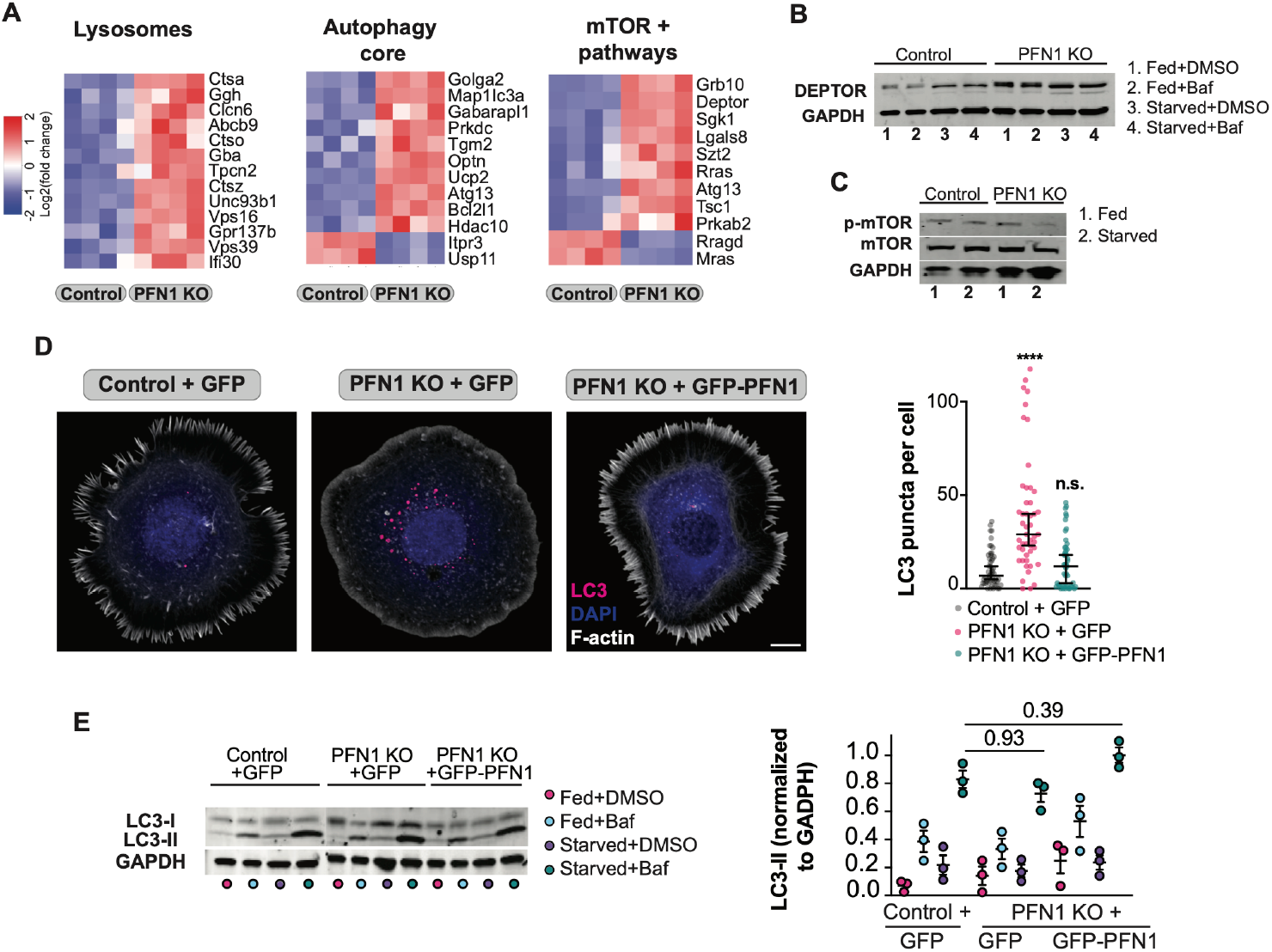
Loss of PFN1 causes an upregulation of autophagy through mTOR deactivation. **A)** Heat maps of RNA-seq analysis performed on PFN1 KO cells and controls, showing differential expression of genes involved in autophagy, endosome/lysosome and mTOR signaling pathways (p < 0.05). **B)** Representative DEPTOR Western blot for PFN1 KO and Control cells under fed and starved conditions, with and without Bafilomycin A, a specific inhibitor of autophagosome-lysosome fusion (n = 3). **C)** Representative Western blot of PFN1 KO and Control cells under fed/starved conditions confirming decrease in phosphorylated mTOR (n = 3). **D)** Representative maximum intensity projection images and quantification (graph) of control, PFN1 KO and PFN1 KO-rescued cells expressing mRuby2-LC3, showing increased LC3 puncta in PFN1-KO cells compared to control, which can be rescued with PFN1-GFP (n = 60, 3 biological replicates), scale bar = 10 μm. P-values are reported relative to Control + GFP; **** = p < 0.0001, n.s. = p > 0.05 **E)** Western blot and quantification of autophagic flux showing LC3I and LC3II levels compared to GAPDH in Control, PFN1 KO and PFN1 KO-rescue cells under fed and starved conditions, with and without Bafilomycin A (n = 3). P-values are listed above conditions being compared.

The increase in autophagosomes observed in PFN1 KO cells could be explained by a defect in protein and organelle homeostasis, resulting in decreased turnover, or PFN1 KO cells might have an increased need for autophagy. Therefore, to test if autophagy was still functional without PFN1, we performed an autophagic flux assay. Quantitative western blot analysis measuring the amounts of soluble LC3-I and lipid bound LC3-II was used to determine autophagosome formation after macro-autophagy was induced by nutrient deprivation. Additionally, to measure total autophagosome production, Bafilomycin A1 was used to prevent autophagosome turnover by inhibiting fusion with the lysosome ^24^. As shown in figure 1 E, autophagic flux was comparable in control, PFN1 KO, and PFN1 KO cells rescued with GFP-PFN1, indicating that there was no general deficiency in autophagosome biogenesis or turnover (Fig. 1E).

### Autophagy induced by the loss of PFN1 selectively targets mitochondria

Since macro-autophagy appeared to be functional in PFN1 KO cells, we next investigated whether a selective form of autophagy was being induced. A clue came from the RNA-seq data, where we found that several major modulators of mitophagy were differentially expressed upon the loss of PFN1. This included increased expression of *TGM2, OPTN*, and *BCL2L1*, and downregulation of the mitophagy inhibitor *SIAH3*^25-28^ (Fig S1). Also, to confirm activation of autophagy in the absence of PFN1, we performed transmission electron microscopy of control and PFN1 KO cells. The electron micrographs clearly showed abundant double membrane bound autophagic vesicles in the PFN1 KO compared to controls (Fig. 2A, arrow). Moreover, most of the autophagic vesicles in PFN1 KO cells contained mitochondria (Fig. 2A, star).

**Figure 2.**
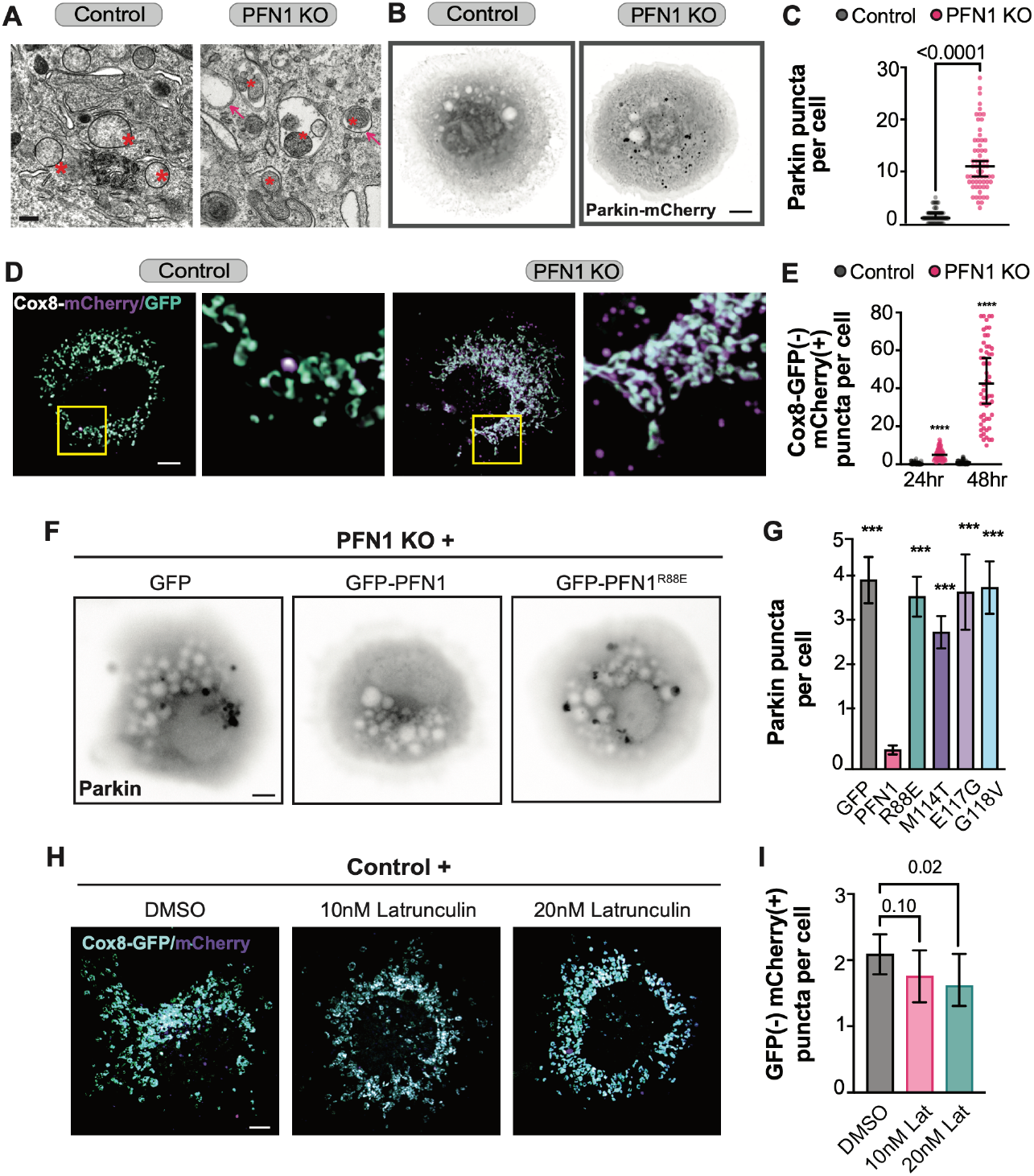
Autophagy induced by the loss of PFN1 selectively targets mitochondria. **A)** Representative electron micrographs of control and PFN1 KO cells. Control cells show healthy mitochondria (stars) and no obvious signs of autophagy, whereas PFN1 KO cells have abundant double membrane bound autophagic vesicles (arrows) many of which contain mitochondria (stars), scale bar = 1 μm. **B-C)** Representative maximum intensity projection images and quantification of control and PFN1 KO cells expressing Parkin-mCherry, showing a 10-fold increase in Parkin foci in PFN1 KO cells compared to controls (n = 60, 3 biological replicates), scale bar = 10 μm. **D-E)** Representative maximum intensity projection images (**D**) and quantification (**E**) of control and PFN1 KO cells expressing the mitophagy reporter Cox8-mCherry-GFP, measured at 24 and 48h post transfection. The amount of mCherry only puncta increased to approximately 10-fold after 48 hr, indicating a massive clearance of mitochondria through mitophagy in PFN1 KO cells compared to controls. P-values are relative to control; **** = p < 0.0001. **F)** Representative epifluorescence images of endogenous Parkin foci in PFN1 KO cells expressing either GFP, GFP-PFN1 or the non-actin binding mutant GFP-PFN1^R88E^. **G)** Quantification of Parkin foci in PFN1 cells expressing GFP, GFP-PFN1 and GFP-PFN1^R88E^ and the ALS associated mutants M114T, E117G and G118V, showing that rescue was only possible with functional PFN1. P-values are relative to PFN1; *** = p < 0.001. **H-**Representative maximum intensity projection images (**H**) and quantification (**I**) of control CAD cells expressing Cox8-mCherry-GFP, treated with low overnight doses of Latrunculin A (10-20 nM) to depolymerize actin roughly 40-50% of total actin to approximate the F-actin loss seen in PFN1 KO cells. Quantification (**I**) shows no significant increase in mitophagy after overnight treatment of Latrunculin A (n = 60, 3 biological replicates). P-values are listed above conditions being compared. Scale bar = 10 μm.

The best described mitophagy pathway involves PTEN-induced kinase 1 (PINK1), which activates the E3 ubiquitin ligase Parkin to tag depolarized and damaged mitochondria for degradation^25-28^. Siah3 inhibits PINK1/Parkin translocation to damaged mitochondria^29,30^, and its downregulation in PFN1 KO (Fig. S1) cells suggested an increase in PINK1/Parkin-based mitophagy. Indeed, when control and PFN1 KO cells were electroporated with a Parkin-mCherry plasmid, we found that PFN1 KO cells had a ten-fold increase in the amount of Parkin puncta (Fig. 2B-C). However, since mitophagy is regularly used by cells to turn over mitochondria; the increase in Parkin puncta could be caused by either an increase in damaged mitochondria or a late-stage defect in mitophagy that would cause mitophagosomes to accumulate. To determine if mitochondria in PFN1 KO cells are successfully degraded by the PINK1/Parkin mitophagy pathway, we used the inner membrane targeted mitophagy reporter Cox8-GFP-mCherry^31^. In acidified compartments, the GFP fluorescence is rapidly quenched but the mCherry fluorescence remains intact. Once mitochondria are delivered to autolysosomes via the fusion of lysosomes with autophagosomes containing mitochondria, the number of reporter puncta with mCherry fluorescence only, increases^31,32^. Using this approach, we found a significant increase in mCherry only puncta in PFN1 KO cells compared to controls (Fig. 2D-E), which further increased to a 10-fold difference after 48 hrs. Therefore, mitochondria are being successfully targeted to lysosomes and degraded in PFN1 KO cells, thus the increase in Parkin foci is likely due to accumulated mitochondrial damage.

Next, we re-expressed PFN1 in PFN1 KO cells to verify that the accumulation of damaged mitochondria was indeed caused by loss of PFN1 expression. Additionally, we expressed the non-actin binding synthetic mutant *R88E*^*20*^ as well as PFN1 mutants causative (*M114T* and *G118V*) or risk-associated (*E117G*) for ALS^33^. PFN1 KO cells expressing the GFP-PFN1 variants described above were assessed for mCherry-Parkin foci formation. Only cells expressing wild-type PFN1 were rescued (Fig. 2F-G). Interestingly, the mutants used for this assay have varying effects on PFN1’s ability to operate as an actin assembly factor, from complete to partial loss of function (Cisterna *et al* 2023, in preparation). Decreasing the amount of polymerized actin in control cells by 40-50% with a low overnight dose of Latrunculin A (10-20 nM) (Cisterna *et al* 2023, in preparation) to approximate the loss of actin caused by PFN1 KO cells, did not result in more mitochondria being delivered to lysosomes (Fig. 2H-I) or cause an increase in the formation of Parkin foci (Fig. S2). Therefore, depolymerizing actin is not by itself sufficient to damage mitochondria and activate mitophagy, indicating that PFN1 has a more specific role in maintaining mitochondrial integrity.

### Loss of PFN1 disrupts mitochondrial metabolism, morphology, and dynamics

Having determined that loss of PFN1 causes upregulation of mitophagy, we next examined the cause of targeted mitochondria degradation. First, we assessed if there was metabolic dysfunction using the Seahorse Extracellular Flux Analyzer to measure glycolysis and oxidative phosphorylation. These assays revealed that PFN1 KO cells have deficient basal and compensatory glycolysis compared to control cells, as well as a slight reduction in basal respiration. (Fig. 3A-F). We also used CellROX Deep Red, a fluorogenic probe for measuring oxidative stress in live cells and found that PFN1 KO cells have a 50% reduction in ROS production compared to control cells (Fig. 3G). Taken together this data shows that loss of PFN1 causes functional defects to mitochondria.

**Figure 3.**
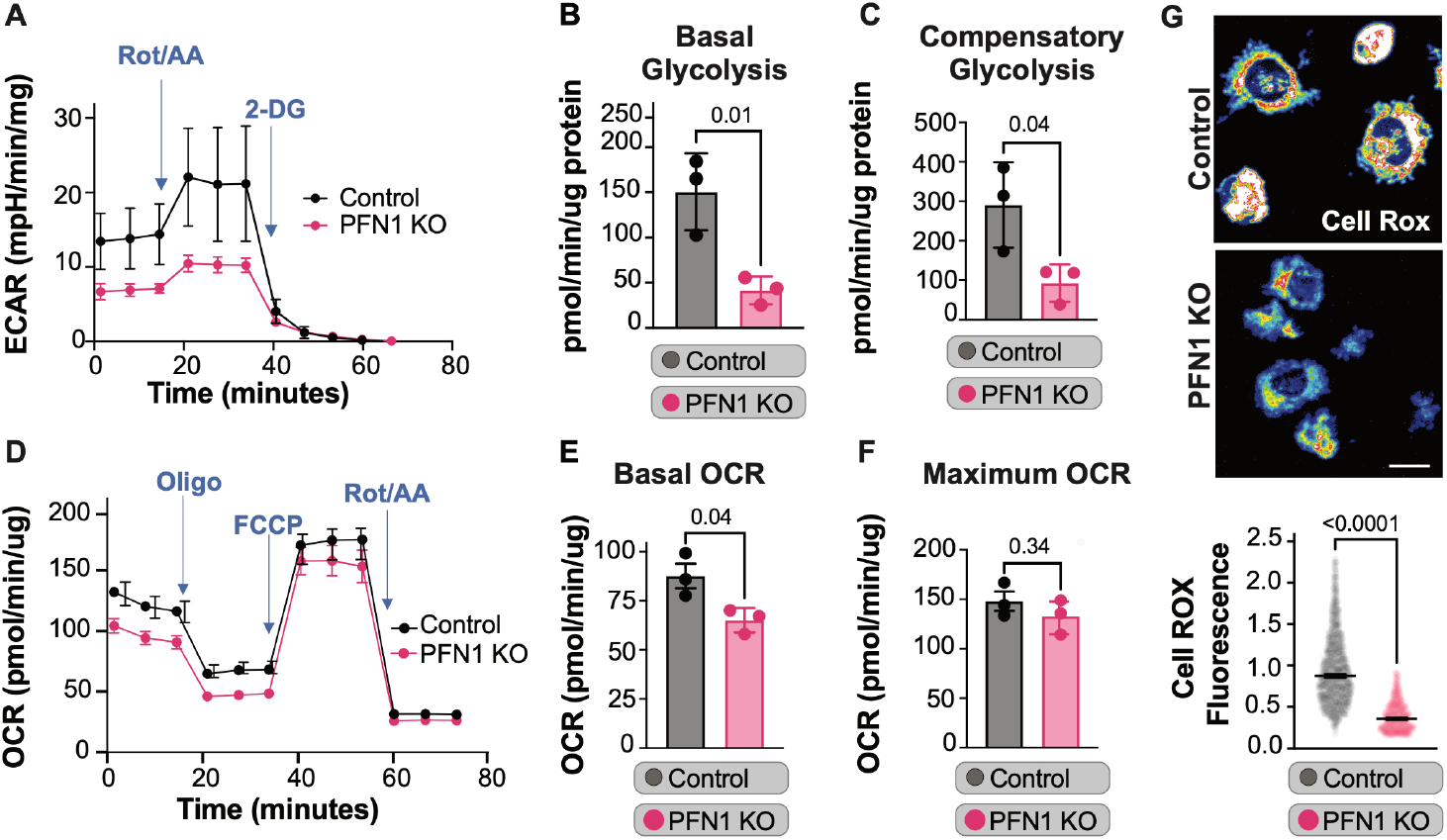
Loss of PFN1 disrupts mitochondrial metabolism. **A-F)** Seahorse Extracellular Flux Analyzer measurements of glycolysis and oxidative phosphorylation. **(A-C)** shows that PFN1 KO cells have deficient basal and compensatory glycolysis compared to control cells, **(D-F)** shows a slight reduction in basal respiration in PFN1 KO cells (n = 3). **G)** Representative images and quantification of control and PFN1 KO cells stained with CellROX Deep Red, a fluorogenic probe for measuring oxidative stress in live cells, scale bar = 40 μm. Quantification shows that PFN1 KO cells have a 50% reduction in ROS production compared to control cells (n = 60, 3 biological replicates). P-values are listed above conditions being compared.

We next investigated whether loss of PFN1 expression affected mitochondria morphology. Transmission electron microscopy images showed distinct differences in mitochondria structure in PFN1 KO cells, namely elongated mitochondrial networks with dysmorphic cristae and many accounts of formation and release of mitochondrial-derived vesicles (MDVs) (Fig. 4A). MDVs are another means of mitochondrial quality control where damaged mitochondrial components can be directed to the cell’s degradative machinery^34-36^. Their presence further confirms that PFN1 deficient mitochondria are damaged and in need of cellular disposal.

**Figure 4.**
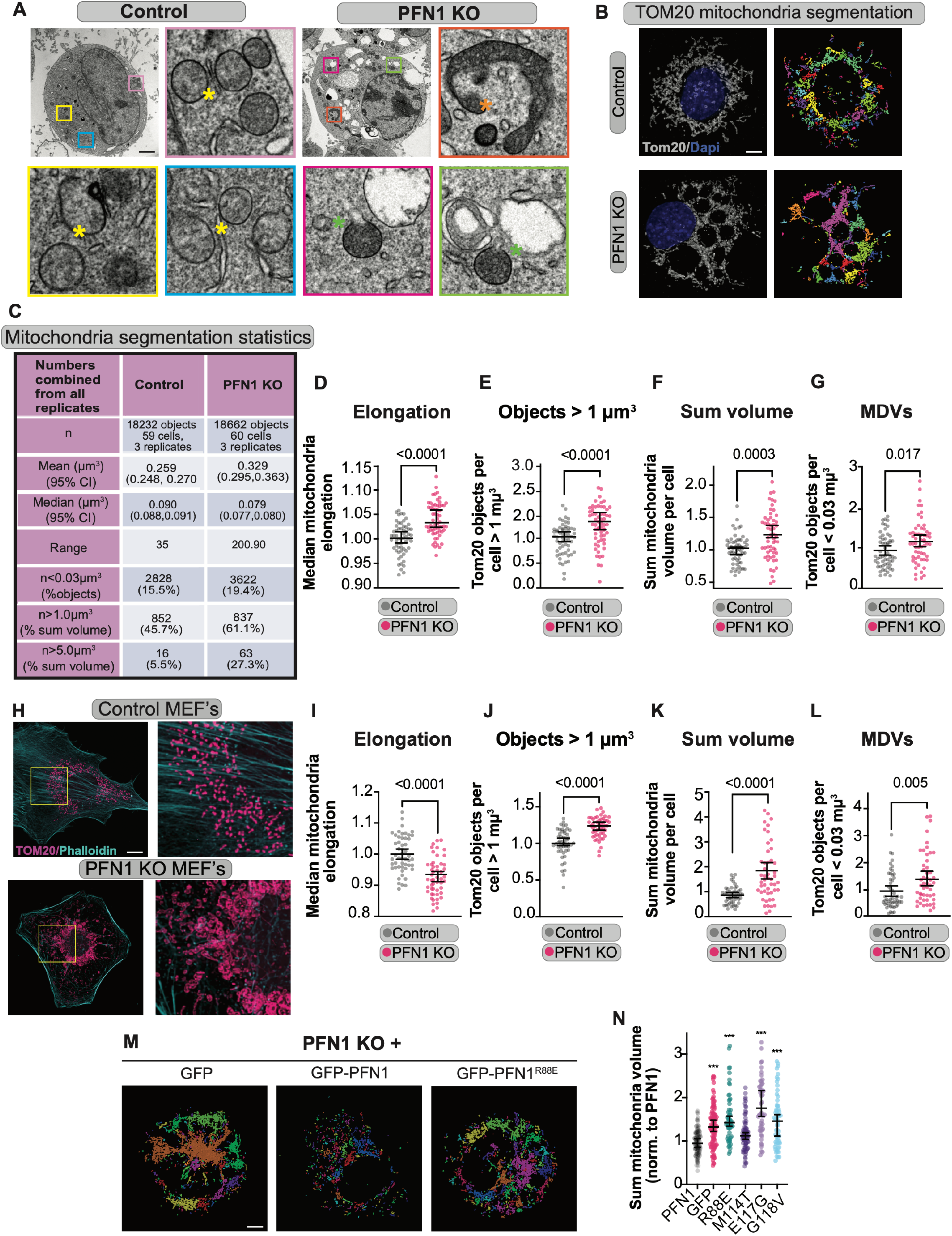
Loss of PFN1 disrupts mitochondrial morphology. **A)** Representative electron micrographs of control and PFN1 KO cells. Control cells have healthy mitochondria (yellow stars) whereas PFN1 KO cells have elongated dysmorphic mitochondria (orange stars) and numerous Mitochondria Derived Vesicles (MDV’s) (green stars), scale bar = 10 μm. **B)** Representative maximum intensity projection images of Tom20 labeled mitochondria from confocal z-stacks in control and PFN1 KO cells and their corresponding segmentation (each binary object is assigned a random color). Scale bar = 10 μm. **C)** Table with mitochondria segmentation statistics. **D-G)** Quantification of volume, size and elongation of Tom 20 labeled mitochondria showing PFN1 KO cells score higher than controls in all parameters measured. The number of Tom20 labeled objects that met the size criteria for mitochondria derived vesicles (MDVs), was also higher in PFN1 KO cells than controls. P-values are listed above conditions being compared. **H)** Representative confocal images showing maximum intensity projections of Tom20 labeled mitochondria in mouse embryonic fibroblast (MEF) control and PFN1 KO cells, scale bar = 20 μm. **I-L**) Quantification of volume, size and elongation of mitochondria and the presence of MVDs showing that, except for elongation, PFN1 KO MEFs have the same mitochondrial phenotypes seen in PFN1 KO CAD cells. P-values are listed above conditions being compared. **M)** Representative images of segmented Tom20 labeled mitochondria in PFN1 KO cells expressing GFP, GFP-PFN1 or the non-actin binding mutant GFP-PFN1^R88E^. Only wild-type PFN1 was able to rescue the increased sum volume phenotype. Scale bar = 10 μm. **N)** Quantification of sum mitochondria in PFN1 KO cells expressing either GFP-PFN1, GFP, GFP-PFN1^R88E^ or the ALS-linked mutations GFP-PFN1^M114T^, GFP-PFN1^E117G^ and GFP-PFN1^G118V^. Complete sum mitochondria volume rescue was only achieved with functional GFP-PFN1 and a partial rescue with the ALS-linked GFP-PFN1^M114T^ mutant. P-values listed are relative to PFN1; *** = p < 0.001.

To quantify these morphological features, we performed immunocytochemistry of the outer mitochondrial membrane (OMM) protein Tom20 and imaged the cells using optical pixel reassignment (SoRa) super-resolution spinning disk confocal microscopy. This was followed by 3D binary segmentation of the z-stack images to obtain the size, shape, and position of all mitochondria present in the cell (Fig.4B-C). PFN1 KO mitochondria were more elongated and present in larger networks than control cells (Fig.4D and E). PFN1 KO cells also had a higher sum volume of mitochondria (Fig. 4F). Additionally, we measured all Tom20-positive mitochondria fragments that met the size criteria for MDVs^34-36^ and found a significantly larger population of MDVs in the PFN1 KO cells (Fig. 4G). These changes in mitochondria morphology were not limited to CAD cells. When we knocked out PFN1 in mouse embryonic fibroblasts (MEFs) we obtained nearly identical results (Fig. 4H-L), with the exception that mitochondria elongation was reduced in PFN1 KO MEFs (Fig. 4K). Rather than being longer, MEF lacking PFN1 had many large, swollen mitochondria that were more circular (Fig. 4H). Re-expressing wild-type PFN1, but not mutants deficient in binding actin (Fig. 4M) or those associated with ALS, rescued all the described morphology phenotypes, including total mitochondria volume (Fig. 4N).

In response to the metabolic demands of the cell, mitochondria divide, fuse, and undergo directed transport^37^. These activities are also necessary for proper mitochondrial function^38^. Additionally, mitochondrial dynamics and mitophagy are closely linked.^39^ To assess mitochondrial dynamics in the absence of PFN1 we performed live cell imaging of cells expressing 4xmts-mNeonGreen to label the inner mitochondrial membrane (IMM)^40^ using optical pixel reassignment spinning disk confocal microscopy. Every mitochondrion in the cell was imaged using a spinning disk confocal microscopy routine that allowed fast volumetric sampling without compromising spatial resolution (detailed in Material and Methods). Mitochondria were then automatically segmented and tracked using MitoMeter software^41^. With this approach, we found that there is only a small reduction in mitochondria velocity and distance traveled in PFN1 KO cells (Fig. 5A-C), indicating almost no change to directed transport. However, there was a significant reduction in the amount of fission and fusion events in PFN1 KO cells compared to controls (Fig. 5D-E). Interestingly, fission and fusion events were also highly correlated to total mitochondria volume in PFN1 KO cells, whereas no correlation was found in control cells (Fig. 5F), suggesting that mitochondria homeostasis is disrupted in the absence of PFN1. Immunolabeling of Drp1, an important fission protein^7^, showed a decrease in Drp1 localization to mitochondria in PFN1 KO cells, validating the live cell data showing decreased fission (Fig. G-H).

**Figure 5.**
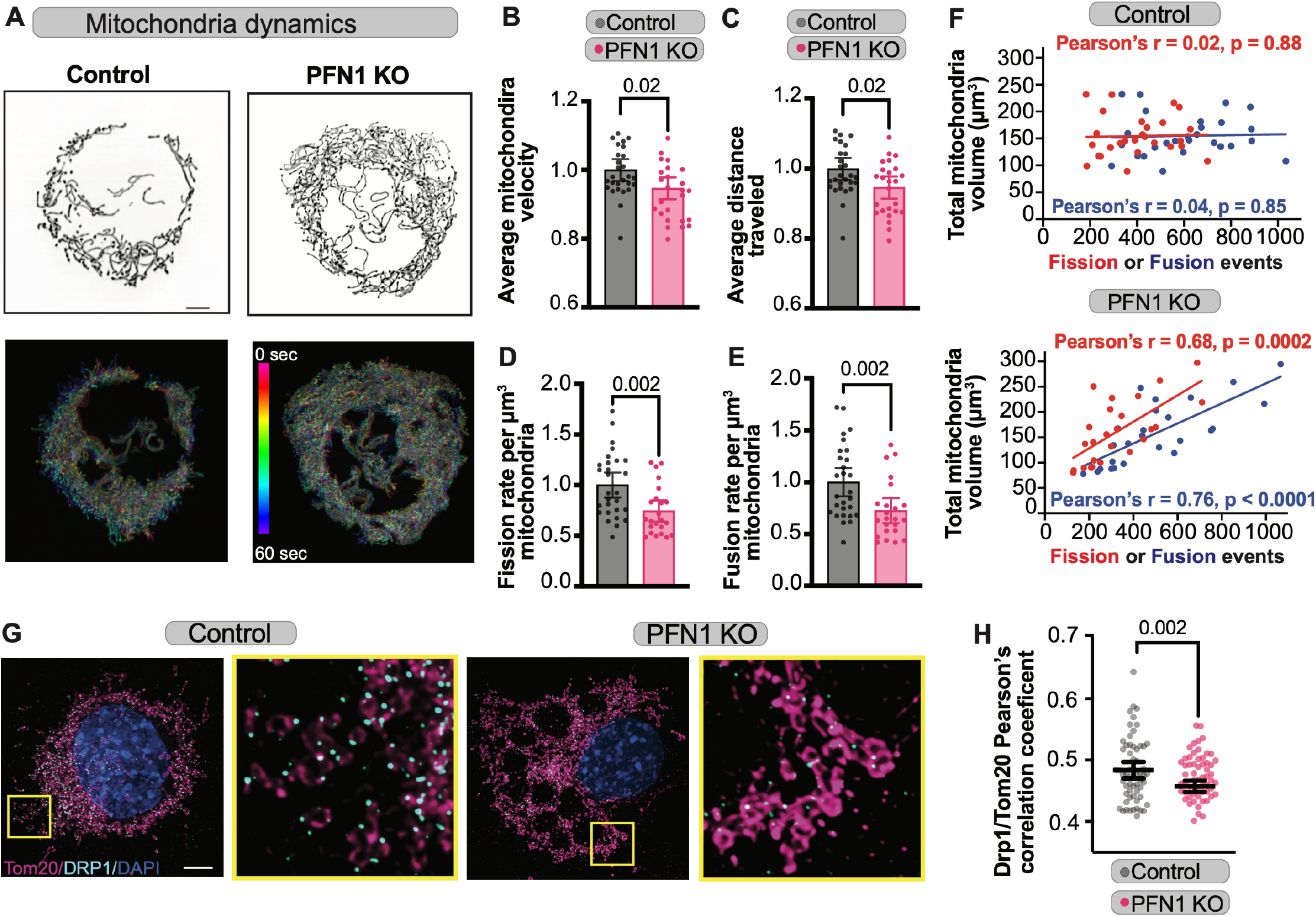
Loss of PFN1 disrupts mitochondrial dynamics. Mitochondrial dynamics were assessed by live cell imaging of control and PFN1 KO cells expressing 4xmts-mNeonGreen to label the inner mitochondrial membrane. **A)** Maximum intensity projections from a single frame (top) and time projections (bottom) of entire movies from representative experiments. Time projections are color-coded according to the scale inserted in right image. Scale bar = 10 μm. **B-C)** Quantification of average velocity and distance traveled of mitochondria in control and PFN1 KO cells, showing no significant differences in these parameters as indicated by p-values above graphs. **D-E)** Quantification of mitochondria fission and fusion events in control and PFN1 KO cells, show significant reductions in both in the absence of PFN1. P-values are listed above the conditions being compared in B-E. **F)** Scatter plots depicting the number of fission or fission events from each experiment plotted against sum mitochondria volume. Linear fitting and calculation of the Pearson’s correlation coefficient (r) reveal that PFN1 KO cells have a high correlation between the number of fission and fusion events with the amount of mitochondria present in the cells, while there is no correlation between these parameters in control cells. Linear fit, Pearson’s correlation coefficient and p-value are reported for each condition; Red for fission and blue for fusion events. **G-H)** Representative images of maximum intensity projections of Tom20 labeled mitochondria colocalized and DRP1 in control and PFN1 KO cells **(G)**, and quantification of the Pearson’s correlation coefficient **(H)**. Colocalization between Tom20 and DRP1 is reduced in the absence of PFN1. P-values are listed above the conditions being compared. Scale bar = 10 μm.

### PFN1 can be found inside the mitochondrial matrix

Having established that PFN1 is necessary for maintaining the functional integrity of mitochondria, we next sought to understand how this is achieved. The most obvious explanation would be that PFN1 facilitates actin polymerization on the mitochondria surface required for fission. However, our rescue experiments showed that even PFN1 mutants that mostly restore actin assembly, could not reverse mitochondrial damage or mitophagy (Fig. 2C and Fig. 4N). Moreover, depolymerizing 40-50% of cellular actin with latrunculin a treatment, does not trigger mitophagy (Fig. 2E). While it could be argued that the specific type of actin filaments involved in mitochondrial dynamics, are not affected in these experiments, another possibility is that PFN1 has some other mitochondrial specific function that has yet to be described, including functions inside the mitochondrial matrix. In fact, both actin and myosin have been identified inside mitochondria and are thought to regulate their function from within^15,42-45^.

Although PFN1 has been shown to localize to non actin structures such as microtubules^46-48^, its interaction with actin networks is transient and therefore not easily observed at specific sites^49^. Additionally, PFN1s high concentration and uniform cytoplasmic distribution further confound colocalization studies^48^. Therefore, to determine if PFN1 was localized to mitochondria, we increased the concentration of detergent in our immunocytochemistry protocol to enhance the post-fixation extraction of cytoplasmic components. With this approach, we found that GFP-PFN1 clearly co-localized with Tom20 labeled mitochondria (Fig. 6A). Additionally, ALS-linked mutations in PFN1 cause the protein to form insoluble aggregates. When expressed in PFN1 KO cells, these mutants would sometimes form aggregates inside of mitochondria, causing them to swell and enlarge (Fig. 6B). This provided additional evidence of GFP-PFN1’s presence inside of mitochondria.

**Figure 6.**
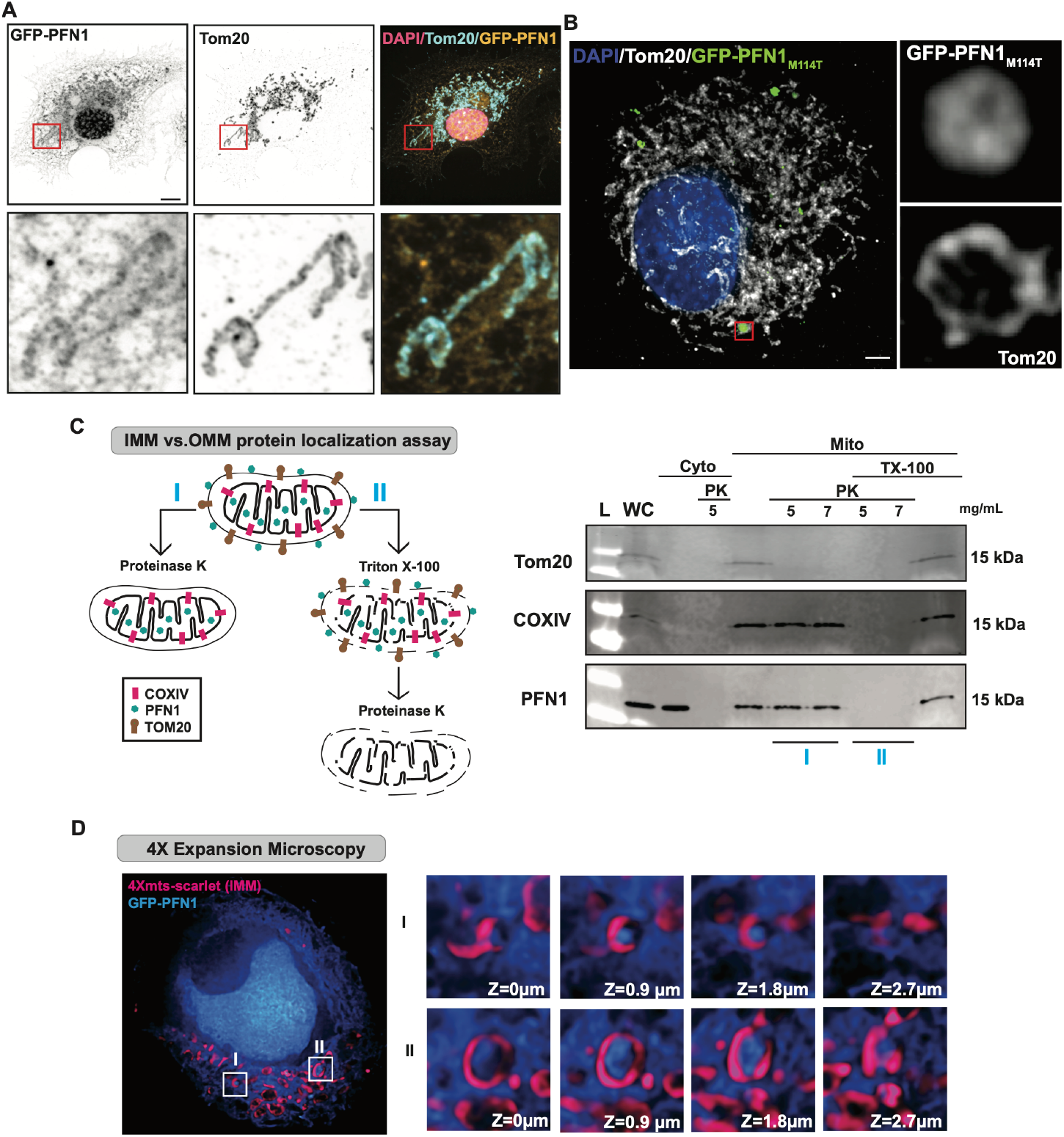
PFN1 is inside the mitochondrial matrix. **A)**Representative images showing maximum intensity projections of Tom20 labeled mitochondria and GFP-PFN1 in PFN1 KO cells after overextraction, showing localization of PFN1 to mitochondria. Scale bar = 10μm. **B)** Representative images showing maximum intensity projections of Tom20 labeled mitochondria colocalized with the aggregating ALS mutant GFP-PFN1^M114T^ in PFN1 KO cells, scale bar = 10 μm. **C)** Schematic protocol and western blot showing the localization of endogenous PFN1 inside the IMM in WT CAD cells. Proteinase K treatment alone removes the outer mitochondria membrane (OMM) protein Tom20 but not the IMM protein COXIV nor PFN1, whereas pretreatment with TritonX-100 removes both the IMM protein COXIV and PFN1. **D)** 4X expansion microscopy of PFN1 KO cells expressing 4Xmts-scarlet to label the inner mitochondria membrane (IMM) and GFP-PFN1, showing that GFP-PFN1 can be visualized inside the IMM.

We next sought to determine if endogenous PFN1 could also be found inside mitochondria and if so, identify which compartment it localized to. We purified mitochondria from CAD cells using the nitrogen cavitation technique, which disrupts the plasma membrane without subjecting mitochondria to shear stress, leaving them fully intact and functional^50^. We divided the mitochondria fraction into two fractions (Fig. 6C). One fraction was treated with Proteinase K to digest all the proteins attached to the outer surface of the mitochondria. Another fraction was treated with Triton X-100 (TX-100) to permeabilize the mitochondria prior to Proteinase K treatment, allowing the digestion of proteins in the inner mitochondrial membrane and the matrix. As shown in Fig. 6C, treatment with Proteinase K alone completely removes the OMM protein TOM 20 from the mitochondria fraction but does not remove the IMM protein COXIV, or PFN1. Pre-treatment with TX-100 followed by Proteinase K successfully removes both the IMM protein COXIV and PFN1 from the mitochondria fraction. These data show that PFN1 is not located on the surface of the OMM, but rather inside the mitochondria membrane.

However, the biochemical analysis of PFN1 at mitochondria fractions did not provide the resolution to determine if PFN1 was inside of the OMM or IMM. With the current resolution of SoRa spinning disk confocal microscopy we could not determine where GFP-PFN1 was localized to in mitochondria. To enhance the spatial resolution and identify if PFN1 was inside the IMM, we performed 4X expansion microscopy^51^ on cells expressing GFP-PFN1 and an IMM localized 4xmts-mScarlet. In z-stacks which captured entire compartments labeled with 4xmts-mScarlet, GFP-PFN1 could be clearly found inside the IMM-labeled boundary (Fig. 6D). Thus, PFN1 resides inside the matrix when localized to mitochondria.

## Discussion

In this study we demonstrated that the actin-monomer binding protein PFN1 is critical for mitochondrial homeostasis. Without PFN1, mitochondria become large and elongated (Fig. 4A, D), have reduced dynamics (Fig. 5D, E) and are defective in glycolysis (Fig. 3B,C). This in turn creates a massive pressure on the cell to increase mitochondria quality control mechanisms such as mitophagy (Fig. 2B, C) and the production of MVDs (Fig. 4A,G). Interestingly, PFN1 is not needed for functional autophagy/mitophagy (Fig. 1D, E and Fig. 2C), and simply decreasing the amount of polymerized actin by itself does not prevent completion of mitophagy by autophagosome-lysosome fusion (Fig. 2E). This was surprising, since dysregulation of autophagy in response to actin depolymerization has been reported by others^52-57^. However, an important difference in our study is that we studied autophagy/mitophagy under conditions where only 40-60% of actin was depolymerized^20^ whereas previous work was done under conditions where nearly all actin filaments were depolymerized. This suggests that the actin filaments needed for autophagosome dynamics, which are a tiny fraction of the total actin within the cell, are not significantly affected when bulk actin assembly is dampened. Interestingly, it is the largest actin monomer consuming structures that are depleted in PNF1 KO cells^20^ and with low concentration Latrunculin A treatment, suggesting that these specific structures are dispensable for autophagy/mitophagy.

Here, we have identified a form of mitophagy that is independent of profilin-actin yet triggered by PFN1 loss of function. As mentioned above, mitophagic flux is functional in PFN1 KO cells (Fig. 2C), and in control CAD cells during drug-induced partial depletion of actin filaments (Fig. 2F), therefore the upregulation of autophagy/mitophagy is likely triggered by chronic mitochondrial dysfunction (Fig.3-5). In fact, despite the upregulation of multiple quality control pathways, dysfunctional, dysmorphic mitochondria still accumulate in PFN1 KO cells (Fig. 2, 4). This was interesting since there has been compelling evidence linking actin binding proteins to mitophagy. For example, cofilin potentiates PINK1/PARK2-dependent mitophagy^58^ and MYO6 (myosin VI) forms a complex with PARK2 and initiates the assembly of F-actin cages to encapsulate damaged mitochondria^59,60^. Additionally, mitophagy requires fission of the damaged portion of the organelle, and several fission pathways are known to require polymerization of actin on the mitochondrial surface^6,61^. This includes fission involving the formin INF2, which requires profilin-actin to assemble filaments^4,5,62^. Although we found that mitochondria fission and fusion were significantly reduced in the absence of PFN1, the overall effect was small and not sufficient to explain the vast mitochondria defects found in PFN1 KO cells. Furthermore, the activated mitophagy phenotype could only be rescued with wild-type PFN1 and not with the non-actin binding PFN1 mutant R88E or the ALS-associated mutants M114T,E117G and G118V (Fig. 4M, N). This is of particular interest since the E117G mutation does not significantly affect PFN1’s ability to stimulate actin polymerization (Cisterna *et al*. in preparation). While it is possible that these mutations perturb only a specific type of actin assembly, an alternate explanation might be that PFN1 has a less obvious role in regulating mitochondrial homeostasis than that of simply assembling filaments on the mitochondria surface.

In support of this idea, we found the PFN1 localizes specifically to the mitochondria matrix (Fig. 6D). Since this is a highly gated cellular compartment,^63,64^ it is extremely unlikely that PFN1 would be there by coincidence. Interestingly, a few studies have found that actin and actin binding proteins localize inside of the mitochondria^42-45^, though none of these experiments determine if localization is within the IMM, as we have shown for PFN1 (Fig. 6D). Functional studies have implied that actin localization to mitochondria is important for maintaining their morphology and respiratory properties^44^. It has yet to be determined what role PFN1 plays within the mitochondrial matrix, given the difficulty of separating its function in the rest of the cell from its potential activities inside of mitochondria. Challenges include separating out indirect effects on mitochondria by proteins that bind to actin filaments in the cytosol^65,66^, or by actin polymerization at the mitochondria surface, which is also linked to mitochondrial respiration^67^. However, it has long been speculated that there are cytoskeletal elements within mitochondria that provide additional forces needed for dynamics, particularly since swelling agents have non-uniform effects on mitochondria morphology (recently reviewed by Zorov *et al*.^68^). Could PFN1’s profound effects on mitochondria homeostasis be explained by its activities within the matrix? We also found the ALS-linked mutants of PFN1 tend to aggregate in mitochondria and cause them to deform (Fig. 6C). This gain of toxic function, combined with loss of function effects on mitochondria morphology and function, could explain how mutant PFN1 causes cellular toxicity and drives neurodegeneration in ALS.

## Materials and Methods

### Cell lines and cell culture

#### Cath. A differentiated (CAD)

cells Control and PFN1 KO cells were generated from CAD cells^69^ originally purchased from Sigma-Aldrich as previously described^20^. CAD cells were cultured in DMEM/F12 medium with L-Glutamine (Gibco) supplemented with 8% fetal calf serum (R&D Systems) and 1% penicillin-streptomycin (Corning). Prior to imaging, CAD cells were plated onto coverslips that had been coated with 10 μg/mL laminin from Engelbreth-Holm-Swarm murine sarcoma basement membrane (Sigma-Aldrich) overnight at 4°C. DMEM/F12 medium without phenol red (Gibco) supplemented with 15mM HEPES (MP Biomedical) was used for live-cell imaging. To assess autophagic responses, we cultured cells for 6 hours in serum- and amino acid-free medium (Krebs–Henseleit medium) which initiates autophagy by starvation. CAD cells differentiate into a neuronal-like cell morphology upon serum withdrawal, but only after 48 hours^69^. We routinely use serum withdrawal to validate CAD cells by ensuring that they can undergo neuronal differentiation as evidenced by the formation of long (> 100 μm), narrow projections after 2 days. PFN1 KO cells were regularly checked for PFN1 expression by western blot.

#### Mouse embryonic fibroblasts (MEFs)

PFN1 KO MEF clonal lines were established from previously described ARPC2 conditional knockout mice^70^. Among these clonal lines, JR20s were used to express Cas9 and our previously verified sgRNA (5’-TCGACAGCCTTATGGCGGAC-3’) targeting mouse PFN1^20^ from pLentiCRISPRv2 (Addgene #52961; a gift from Feng Zhang) via lentiviral transduction. Lentivirus was generated through the transfection of pCMV-V-SVG, pRSV-REV, pMDLg/pRRE, and pLentiCRISPRv2-PFN1sgRNA (500 ng each) into HEK293FT cells using X-tremeGENE HP transfection reagent (Roche). Lentivirus was harvested 72 hours later and subsequently used to infect JR20 cells supplemented with 4 μg/mL of Polybrene. Roughly 72 hours following the addition of lentivirus, JR20s expressing PFN1 sgRNA and Cas9 were selected using 2 μg/mL puromycin treatment for 48 hours. A second round of infections was used for the lentiviral expression of Cox8-GFP from a pLentiLox5.0 vector in a similar manner as described above (to label mitochondria). MEF cells were cultured in DMEM medium with 4.5 g/L D-Glucose, L-glutamine, 25mM HEPES, and without Sodium Pyruvate (Gibco) supplemented with 8% fetal calf serum (R&D Systems) and 1% penicillin-streptomycin (Corning). Prior to imaging, MEFs were plated onto coverslips that had been coated with 10 μg/mL human fibronectin (Corning) for 1 hr. at room temperature (RT). All cell lines used for this study were routinely tested for mycoplasma using the Universal Detection Kit (ATCC).

### DNA constructs

The following DNA constructs purchased from Addgene were used: EGFP-C1(Plasmid #54759,), mEGFP-PFN1 (Plasmid #56438), Cox8EGFP-mCherry (Plasmid # 78520), 4xmts-mScarlet-I (Plasmid #98818), 4xmts-mNeonGreen (Plasmid #98876), pLentiCRISPRv2 (Plasmid #52961) mCherry-Parkin (Plasmid #23956). GFP-PFN1 and GFP-PFN1^R88E^ have been previously described^71^. Plasmids expressing the PFN1-ALS mutants *M114T, E117G* and *G118V* were generated from GFP-PFN1 plasmid using site-directed mutagenesis (Q5 New England Biolabs) with the following primers: M114T: gtc ctg ctg acg ggc aaa gaa g (forward) and cc ttc ttt gcc cgt cag cag gac (reverse); E117G: atg ggc aaa gga ggt gtc cac (forward) and g gac acc tcc ttt gcc cat c (reverse); G118V: atg ggc aaa gaa gtt gtc cac ggt ggt ttg (forward) and caa acc acc gtg gac aac ttc ttt gcc cat (reverse). Mutagenesis was confirmed by sequencing (Genewiz). mRuby2-LC3 was generated by subcloning LC3 from pmRFP-LC3 (Plasmid #21075) into the pcDNA3-mRuby2 (Plasmid # 40260) using the 5′ BamHI and the 3′ EcoRI cloning sites. Correct inserts were confirmed by sequencing (Genewiz). All constructs were prepared for transfection using either the GenElute HP Endotoxin-Free Plasmid Maxiprep Kit (Sigma-Aldrich) or NucleoBond Xtra Midi EF kit (MACHEREY-NAGEL).

### DNA electroporation

The Neon Transfection System (Invitrogen) was used to introduce DNA constructs into cells using the 10 μL transfection kit^71^. Briefly, cells were grown to of 70-80% confluency, detached using 0.5% Trypsin/EDTA solution (Corning) and pelleted by brief centrifugation. Pellets were rinsed with Dulbecco’s Phosphate-Buffered Saline (DPBS, Corning) and resuspended in a minimum amount of buffer R (Invitrogen) with a total of 1μg of DNA per electroporation reaction. Thereafter cells were cultured for 14-48 hours in antibiotic-free medium prior to additional experimental procedures.

### LC3, DEPTOR and mTOR western blots

Adherent cells were harvested in RIPA buffer with cOmplete EDTA-free Protease Inhibitor Cocktail Roche (Millipore Sigma), except for probing for p-mTOR, where phosphatase inhibitor (Roche Phospho-Stop, Sigma) was also added. Whole cell lysates were prepared by membrane disruption using repeated passage through a 27 gauge needle. Protein content was assessed with Pierce BCA Protein Assay Kit (Thermo Fisher) and diluted in SDS buffer stained with Orange G (40% glycerol, 6% SDS, 300 mM Tris HCl, pH 6.8). 10 μg samples were evenly loaded on an SDS-PAGE gel (Novex 4-20% Tris-Glycine Mini Gels, Thermo Fisher, or 15% gel as indicated), except for p-mTOR where 25ug was added. Protein was transferred to a PVDF membrane (0.2 micron, 0.45 micron for mTOR and p-mTOR, both from Immobilon) and blocked in 5% Bovine Serum Albumin (BSA) (Sigma-Aldrich) for 20 mins. All antibodies were diluted in 5% BSA and 0.1% Tween-20 (Fisher Scientific). Primary antibodies were incubated at 4°C overnight and secondary antibodies (Li-Cor; Abcam) were incubated for 2 hours at RT. LC3 and GAPDH from whole cell lysate were detected with Li-Cor fluorescent antibodies on an Odyssey detection system (Li-Cor) or via X-ray film after incubation with a developing reagent (Thermo Fisher), as indicated. WesternSure Pre-Stained Chemiluminescent Protein Ladder (Li-Cor) was used as a molecular weight marker. The following antibodies/dilutions were used: rabbit anti-GAPDH (cat # 2118, 1:3000 dilution, Cell Signaling Technology), rabbit anti-LC3 (cat # 2775, Cell Signaling), rabbit anti-DEPTOR/DEPDC6 (cat # NBP1-49674, Novus biologicals), rabbit anti-mTOR (7C10) (cat # 2983, Cell Signaling) and rabbit-Phospho-mTOR (Ser2481) (cat # 2974, Cell Signaling). For secondary antibodies, goat anti-rabbit Alexa Fluor™ 680 (Li-Cor) was used at 1:3500 dilution for imaging on the Li-Cor Odyssey detection system and goat anti-rabbit HRP (Abcam) was used for X-ray detection. For quantitative westerns, antibody detection was determined to be in the linear range by loading increasing lysate concentration as a function of signal.

### Oxygen Consumption and Glycolytic Rates Measurement

The Seahorse XFe96 Extracellular Flux Analyzer was used to measure oxygen consumption rates (OCR) and glycolytic rates of adherent CAD cells in real time. A range of cell seeding densities and a series of titration experiments were initially tested to determine optimal conditions. One day prior to assay, the Seahorse Sensor Cartridge (#102416-100, Agilent) was hydrated in Seahorse XF Calibrant (#102416-100, Agilent) overnight at 37 °C in a non-CO2 incubator. Utilizing an XF 96 cell culture microplate (#102416-100, Agilent), 1x10 ^4^ CAD cells were plated and maintained in appropriate growth medium. On the day of the assay, cells were examined under the microscope to confirm confluence. Growth medium was gently aspirated, and cells were washed two times with freshly prepared Seahorse XF assay medium (#103680-100, Agilent) supplemented with glucose (10mM), pyruvate (1mM), and glutamine (2mM). The cell culture microplate containing the cells was incubated at 37°C in a non-CO2 incubator for 45-60 minutes, and then transferred to Seahorse XFe96 Extracellular Flux Analyzer (# S7894-10000, Agilent). The sensor cartridge and cell plate were equilibrated and calibrated according to the manufacturer’s instructions. To measure OCR, the cells were then subjected to Mito Stress Test (#103015-100, Agilent) by sequential injections of mitochondrial inhibitors to constitute final concentration per well of 1μM Oligomycin, 1μM Carbonyl cyanide-4 (trifluoromethoxy) phenylhydrazone (FCCP), and 0.5μM Rotenone/Antimycin A. All dilutions were freshly reconstituted the same day. Analysis was run using the standard Mito Stress assay protocol. Oxygen consumption rates were determined by Seahorse X96 Wave Software, and the data was normalized to protein concentration per well (pmol/min/μg). To measure the glycolytic rate, Seahorse XF Glycolysis Rate Assay Kit (#103344-100, Agilent) was used. The same protocol for hydrating cartridge, seeding, and washing cells were used as mentioned above. The glycolytic rate was measured by sequential injections of 0.5μM Rotenone/Antimycin A and 50mM of 2-DG. Analysis was run using the standard glycolytic rate assay protocol. Glycolytic rate was determined by Seahorse X96 Wave Software and the data was normalized to protein concentration per well (pmol/min/μg).

### Microscopy

Unless noted, all images were acquired using either a Nikon A1R+ laser scanning confocal microscope or a Nikon CSU-W1 SoRa spinning disk confocal microscope. The A1R+ microscope was equipped with a 1.49 NA 60X Apo TIRF objective, GaAsP detectors, and a piezo stage. To achieve super-resolution using the laser-scanning confocal, zoom settings were used to obtain confocal z-stacks containing images with oversampled pixels (0.03 μm). Subsequent 3D deconvolution (NIS-Elements) resulted in images with approximately 150 nm resolution. The CSU-W1 SoRa spinning disk confocal microscope was equipped with a 100X 1.49NA SR objective, a 60X 1.49 NA Apo TIRF objective, a 40X 1.25 NA SIL silicone oil objective, a 20X 0.75 NA Plan Apo objective, a piezo stage, a Tokai Hit stage top incubator, and a Hamamatsu Fusion BT camera. Images were acquired using either the W1 or SoRa spinning disk modes depending on the needs of the experiment. To achieve super-resolution, confocal z-stacks were acquired using the 100X objective and the SoRa 2.8X magnifier, followed by 3D deconvolution (NIS-Elements). Images of Parkin foci (figure 2D) were acquired using an EVOS M5000 desktop microscope (Life Technologies) with an Olympus 40X PlanXApo 0.95 NA objective.

### Immunofluorescence

Cells were seeded onto coverslips and cultured for 2 hr prior to fixing and staining. Thereafter cells were fixed with 4% electron microscopy grade paraformaldehyde (Electron Microscopy Sciences) for 10 min at RT and then permeabilized for 3 minutes with 0.2% Triton-X 100 (Millipore Sigma). Over-extracted cells were treated with 0.4% Triton-X 100 for 10 min. Cells were then washed three times with DPBS and stained overnight at 4°C with primary antibodies diluted in PBS. Next, they were washed twice with PBS for 5 min, incubated with secondary antibodies (diluted 1:1000) for 2 hr at RT in PBS. Actin filaments were stained with Alexa Fluor 488, 568 or 647 phalloidin (Life Technologies) for 30 min at RT in DPBS. Cells were washed three times with DPBS before mounting with Prolong Diamond (Life Technologies). The following antibodies were used: Rabbit anti-Tom 20 (cat# 11802-1-AP, Proteintech) 1:300 dilution; rabbit-anti-parkin (cat # PA5-13399, Invitrogen) 1:50 dilution, mouse anti-DLP1 (cat# 611113 Clone 8/DLP1 (RUO), BD biosciences) 1:100 dilution, Preabsorbed secondary antibodies used were anti-mouse IgG 647, anti-rabbit IgG 568 at a 1:1000 dilution. Coverslips were mounted onto slides using ProLong Diamond (Thermo Fisher). All slides were cured at RT in the dark for 2 days prior to imaging.

### Quantification of autophagy and mitophagy

#### Quantification of LC3 puncta

CAD cells were electroporated with a plasmid expressing mRuby2-LC3. The next day, they were plated onto laminin-coated coverslips for 2 hrs, switched to nutrient-deprivation media (Krebs-Henseleit) for 6 hrs, and then fixed as described above. Confocal z-stacks of mRuby2-LC3 were taken using the Nikon A1R+ laser scanning confocal microscope using the 60X objective at settings that met Nyquist criteria. The z-stacks were compressed into a maximum intensity projection, the background was subtracted, and then mRuby2-LC3 was segmented using intensity-based thresholding using size filters (objects needed to be > 0.01 μm^2^). The threshold value was set manually from cell to cell. LC3 puncta were measured both as number of puncta per cell and total LC3 puncta volume, both gave equal results.

#### Quantification of Parkin puncta

Images were taken from at least 20 random fields of view and experiments were performed in triplicate, resulting in at least 60 images for each condition to be used for analysis and quantification. Parkin puncta were counted manually using the following criteria: puncta fluorescence must be twice that of background fluorescence, only single nucleus cells were counted, and cells must have limited direct contact with surrounding cells.

#### Quantification of Cox8-EGFP-mCherry

Cox8 is an inner mitochondrial membrane protein. Therefore, the tandem Cox8-EGFP-mCherry can be specifically targeted to mitochondria and has recently been used to monitor mitophagy in cultured cells. With this construct, normal healthy mitochondria express both EGFP and mCherry whereas autolysosome-enwrapped mitochondria will only display mCherry fluorescence. Therefore, to monitor mitophagy by quantifying mitochondria within autolysosomes, we used the Nikon CSU-W1 SoRa spinning disk confocal microscope 100x objective to image cells, create maximum projection images from z stacks and manually counted mCherry puncta. At least 20 random fields of view were selected for each condition and experiments were performed in triplicate. Puncta selection criteria were the same as for Parkin puncta in addition to picking puncta that expressed mCherry only.

### Reactive oxygen species staining

Control and knock-out cells, plated on laminin coated coverslips were stained for 30 min at 37C with CellROX reagent (Invitrogen) at 1uM final concentration, washed 3 X 5 minutes in DPBS, before being mounted in live cell imaging chambers and imaged at 40X.

### Determining the effect of actin depolymerization on mitophagy

To determine the effects of depolymerized actin on mitophagy, we electroporated WT CAD cells with the Cox8 EGFP-mCherry plasmid, cultured cells on laminin coated coverslips for 48 hours before treating them with either 10nM or 20nM Latrunculin dissolved in DMSO (Sigma). We have previously shown that low dose overnight treatment with Latrunculin A (10-30 nM), reduce the amount of polymerized actin by about 40%, without comprising cell morphology or survival (Cisterna *et al*., 2023). Cells were treated for 14h at 37°C in a standard tissue culture incubator, before proceeding with phalloidin staining to visualize actin and mounting the coverslips for imaging and analysis.

### Electron microscopy

Control and PFN1 KO CAD cell pellets were fixed in 4% paraformaldehyde, 2% glutaraldehyde in 0.1 M sodium cacodylate (NaCac) buffer, pH 7.4, postfixed in 2% osmium tetroxide in NaCac, stained en bloc with 2% uranyl acetate, dehydrated with a graded ethanol series and embedded in Epon-Araldite resin. Thin sections were cut with a diamond knife on a Leica EM UC7 ultramicrotome (Leica Microsystems, Inc, Bannockburn, IL), collected on copper grids and stained with uranyl acetate and lead citrate. Cells were observed in a JEM-1400Flash transmission electron microscope (JEOL USA Inc., Peabody, MA) at 120 kV and imaged with a Gatan “OneView” CCD Camera using DigitalMicrograph software (Gatan Inc., Pleasanton, CA).

### Quantification of mitochondria morphology

Cells were plated onto laminin-coated coverslips for 90 minutes, fixed and stained overnight for Tom20 to label mitochondria. For rescue experiments, cells were transfected 24 hrs prior. Cells were imaged using the Nikon CSU-W1 SoRa spinning disk confocal microscope with a 100X 1.49NA SR using the 2.8X SoRa magnifier. Confocal z-stacks were deconvolved using the Blind algorithm (Nikon Elements) at 20 iterations. This combination of algorithm and number of iterations was chosen because it yielded the highest signal to noise without being destructive to the morphology of Tom20-stained mitochondria. 3D binary objects were created from the deconvolved z-stacks using intensity-based thresholding. The minimum threshold intensity value was manually chosen for each image, and a clean filter was applied to remove punctate background immunofluorescence from the 3D binary mask. Object parameters (volume and elongation) were measured in Nikon Elements software and data was exported into Microsoft Excel for analysis. 2-3 biological replicates were performed per condition.

### Live cell imaging of mitochondria dynamics

#### Live cell microscopy and image processing

Live imaging of mitochondria was performed using the Nikon CSU-W1 SoRa spinning disk confocal microscope. Confocal z-stacks were made using the 40X SIL silicon oil objective, the 4.0X SoRa magnifier, and 2 × 2 camera binning. Image exposure times were ≤ 10 ms so that entire cell volumes could be imaged every 2-5 seconds. Background noise was removed using Denoise.ai (NIS-Elements), a trained neural network that uses deep learning to estimate and remove the noise component of an image. Images were then deconvolved in NIS-Elements using the 3D Blind Deconvolution algorithm. Before analysis, images were exported into Fiji/ImageJ and corrected for photobleaching using the histogram matching algorithm.

#### Quantification of mitochondria dynamics

Analysis of mitochondria dynamics from live cell imaging experiments was performed using a modified version of the Mitometer pipeline^41^. The segmentation of images was achieved by employing a Frangi filter to enhance the visualization of mitochondria^72^ a similar approach as MitoGraph^73^. During frangi filtration, the hessian matrix was thresholded at the square root of the maximum of the frobenius norm for computational efficiency. Semantic segmentation was achieved via a threshold of 1e-05, where anything in the frangi-filtered image above the threshold was kept as a mitochondrial pixel. Subsequently, instance segmentation of individual mitochondria was carried out using connected components derived from the semantically segmented image. Skeletonization of the image was performed using Lee’s method^74^ implemented via scikit-image. Instance segmentation of mitochondrial branches was then carried out by traversal between branch junctions and branch tips, or branch tips and branch tips. Mitochondrial tracking was performed as detailed in Mitometer. Estimates of fission and fusion events were assigned in a binary manner frame to frame by calculating mitochondrial branch number differences between adjacent frames on a mitochondrion-by-mitochondrion basis. All parameters were kept constant for analysis of all images to ensure comparable results between samples.

### Expansion Microscopy

Sample preparation for 4X expansion microscopy was performed using methods and reagents previously published^75^, with some minor adjustments. Cells were electroporated with PFN1-GFP and 4xmts-mScarlet-I (1ug DNA total) and cultured overnight. The next day, cells were seeded onto 13-mm-diameter round coverslips (cat. # 174950, Thermo Fisher,) pre-coated with 10 μg/mL laminin (Sigma) and allowed to adhere for 4h at 37°C in a standard tissue culture incubator. Cells were then fixed with 4% paraformaldehyde (Electron Microscopy Sciences) at RT for 30 minutes, washed with DPBS (Corning), and post-fixed with 0.25% Glutaraldehyde (Electron Microscopy Sciences) for 10 minutes followed by a ten-minute quenching step (100 mM Glycine, Sigma). Next, cells were washed with DPBS and overlaid with 250 μL of gelation solution^75^ and placed on ice for 5 minutes. Thereafter, coverslips with cells were removed and placed on prepared spacer slides^75^ with cells facing downwards. 40 μL of gelation solution was then pipetted between the spacer slides and coverslip and allow to gel completely in a humidity chamber at 37°C for 1 hour. Once the gel had formed, coverslips with gels were removed from the chamber and cooled at RT for 2 min. Next, coverslips with gels were placed in a 10 cm tissue culture dish (Corning), gel facing up, covered with aluminum foil, and incubated for 3 hr at RT in Proteinase K digestion buffer^75^ on an orbital rocker at 60 rpm. After digestion, gels which had now detached from coverslips were removed and placed in a clean 10 cm dish, and expanded by 3 serial incubations in H^2^O on an orbital rocker for 60 minutes at RT. Following complete expansion, the gels were transferred to a 6 cm glass bottom dish pre-coated with 0.1% (weight/volume) Poly-D-lysine (Sigma) for imaging.

### Assay to determine protein localization to the mitochondrial matrix

#### Purification of crude mitochondrial fractions

Adherent cells were grown to 100% confluence in DMEM/F12 medium supplemented with 10% fetal bovine serum and 1% penicillin-streptomycin. Cells were washed twice with DPBS and harvested in Mitochondria Isolation Buffer (MIB) (200 mM sucrose, 10 mM Tris-MOPS, 1 mM EGTA-Tris). Next, cells in suspension were held at 150 psi for 20 min on ice in Pierce Protease Inhibitor Cocktail (Thermo Scientific) in a 45 mL Cell Disruption Vessel (Parr Instrument) with a mini magnetic stirrer^76^. Cells were disrupted by releasing the pressure. Next, the expelled homogenate was palleted by centrifugation for 3 min (at 300 g at 4°C) (Eppendorf). The pellet was discarded, and the supernatant was further separated by centrifugation at 8,000 g for 15 min at 4°C in a microcentrifuge. The pellet which contained the crude mitochondrial fraction (Mito) was retained, and the supernatant was removed and centrifuged at 20,000 g for 20 min at 4°C in a microcentrifuge. The pellet from this final centrifugation step was discarded, and the supernatant containing the Cytosolic fraction (Cyto) was retained.

#### Proteinase K/Triton X-100 treatment

The isolated Mito fraction was incubated with 0 or 1% (v/v) Triton X-100 on ice for 3 min. Then, the Mito and Cyto fractions were both incubated with 0, 0.5, 1, 3, 5, or 7 mg/mL proteinase K (Thermo Scientific) dissolved in MIB for 1 hour at 37°C. Next, proteinase K digestions were stopped by 10 mM phenylmethanesulfonyl fluoride (final concentration) (Protease Inhibitor Cocktail, Promega) on ice for 20 min. The samples were then resuspended in RIPA buffer solution (50 mM Tris-HCl pH 7.4, 150 mM NaCl, 1% (v/v) Triton X-100, 0.5% (w/v) sodium deoxycholate, 0.1% (w/v) sodium dodecyl sulfate) with Protease Inhibitor Cocktail (Promega)^77^. Mito and Cyto samples were finally analyzed by western blot.

#### Western blotting

Whole-cell (WC) lysates were prepared by harvesting with a scraper in RIPA buffer and using repeated passages (5x) through a 21-, 25-, and 27-gauge needle. Then, protein quantification was assessed with Pierce BCA Protein Assay Kit (Thermo Scientific) and diluted in SDS buffer stained with Orange G (40% glycerol, 6% SDS, 300 mM Tris HCl, 5% β-mercaptoethanol pH 6.8). Next, the samples were denatured at 95°C for 5 min before loading. 10 μg of samples were loaded on SDS-PAGE gel (Novex 4%–20% Tris-Glycine Mini Gels, Thermo Scientific). Protein was transferred to a PVDF membrane 0.2 μm (Amersham) and blocked in 5% Bovine Serum Albumin (BSA) (Sigma-Aldrich) for 20 min. All antibodies were diluted in 5% BSA and 0.1% Tween-20 (Fisher Scientific). The following antibodies/dilutions were used: rabbit anti-COXIV (1:2,000 dilution, overnight at 4°C, ab202554, clone EPR9442(ABC), Abcam); mouse anti-TOMM20 (1:1,000 dilution, 2 hours at 37°C, ab56783, clone 4F3, Abcam); mouse-anti-TOM20 (1:500 dilution, 2 hours at 37°C, H00009804-M01, clone 4F3, Abnova); rabbit anti-profilin 1 (1:1,000 dilution, overnight at 4°C, ab124904, clone EPR6304, Abcam). For secondary antibodies, goat anti-rabbit (1:10,000 dilution, 2 hours at RT, 926-32211, Li-Cor) and goat anti-mouse (1:10,000 dilution, 2 hours at RT, 926-32210, Li-Cor) was used for imaging on the Li-Cor Odyssey detection system.

### Statistical Analysis

All data was tested for normality using the Shapiro-Wilk normality test. If the data assumed a Gaussian distribution, groups were compared using either an unpaired student’s t-test for two conditions or by ordinary one-way ANOVA followed by Tukey’s post-hoc test for comparisons of three or more conditions. If the data failed the normality test, then groups of two were compared using the Mann-Whitney test and groups of three or more were compared using the Kruskall-Wallis test followed by Dunn’s multiple comparisons test. Analysis and graphing of results were performed using Graphpad Prism software.

## Acknowledgements

We would like to thank Aleksandra Zamaro (Ilatovskaya lab, Augusta University) for assistance with early attempts at purifying mitochondria from CAD cells. Research reported in this publication was supported by the Maximizing Investigators’ Research Award from the National Institute of General Medical Sciences of the National Institutes of Health under grant number R35GM137959 to EAV.

