## Supplementary Figures and Video Legend for "The actin binding protein profilin 1 is critical for mitochondria function"

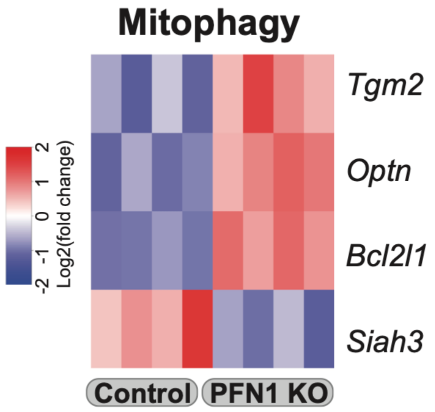


**Figure S1. Loss of PFN1 causes an upregulation of mitophagy.**
Heat map of RNA-seq analysis performed on PFN1 KO cells and controls, showing differential expression of genes involved in mitophagy signaling pathways ($p < 0.05$). Major modulators of mitophagy which were upregulated included *TGM2*, *OPTN*, and *BCL2L1*, whereas of the mitophagy inhibitor *SIAH3* was downregulated.

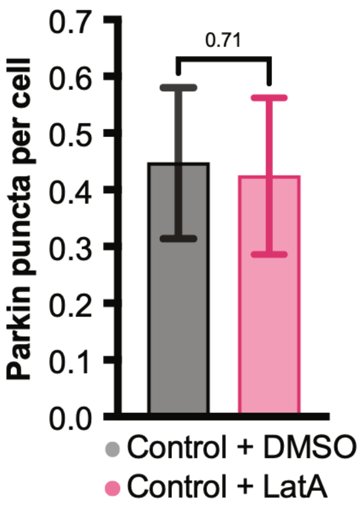


**Figure S2. Actin depolymerization does not by itself induce mitophagy.**
Decreasing the amount of polymerized actin in control cells by 40-50% with a low overnight dose of Latrunculin A (10-20 nM) to approximate the loss of actin caused by PFN1 KO, does not result induce the formation of Parkin foci

### Supplementary Video Legends

**Video S1. PFN1 KO cells have reduced mitochondria dynamics.**
Control and PFN1 KO cells expressing 4xmts-mNeonGreen to label the inner mitochondrial membrane were imaged using optical pixel reassignment spinning disk confocal microscopy. Maximum intensity projections of the confocal z-stacks are shown. Time is indicated in min:sec.
